# Frequency-Dependent Inter-Brain Synchrony is Modulated by Social Interaction in Freely Moving Mice

**DOI:** 10.1101/2024.05.21.593536

**Authors:** Alessandro Scaglione, Jessica Lucchesi, Anna Letizia Allegra Mascaro, Francesco Saverio Pavone

## Abstract

Social interaction, a pivotal aspect of human and animal behavior, involves a dynamic exchange of information that shapes behavioral responses, emotional states, and cognitive processes. To gain insights into the neural mechanisms underlying these processes, it is necessary to simultaneously investigate the brain activity of socially interacting subjects. Commonly, the simultaneous study of behavior and brain activity during the execution of social tasks is conducted through Hyperscanning in humans which limits the availability of interventions. Here we describe a new experimental platform that combines the development of a new miniaturized optical system, the “MiCe-μScope”, to monitor neural activity across the entire cortical mantle with a behavioral paradigm to perform a Hyperscanning study in freely moving mice engaged in social interaction. Our results revealed inter-brain synchrony across different frequency bands widespread over the entire cortical mantle, modulated by social behavior. This finding suggests that synchronization reflects the mutual prediction performed by the entire cortex in mice of interacting dyads. The presence of different synchronization maps in these frequency bands suggests a multiscale nature of interaction, extending the predictive nature of interaction to cortical areas beyond the medial prefrontal cortex. Our work provides an experimental framework to conduct Hyperscanning studies in an animal model that mirrors findings from human studies.

## INTRODUCTION

Social interaction is a fundamental aspect of both human and animal behavior. Individuals participating in social behavior act and react with each other generating a dynamic process of mutual information exchange^1^. This continuous exchange shapes behavioral responses, emotional states and cognitive processes, highlighting the complex nature of social dynamics in human societies and the animal world. The exploration of neural activity, underlying social behavior, has often focused on analyzing the neural circuits of a single animal engaged in interactions with others^2–5^. This approach captures only a partial spectrum of the information that could be collected from recording the neural processes of two or more animals interacting simultaneously. As a solution, the idea of conducting Hyperscanning studies emerged: the simultaneous and real-time measurement of the brain activity of multiple subjects engaged in social interactions, aimed at revealing neuronal inter-brain correlations during different behavioral tasks^6–11^. In contrast to conventional behavioral paradigms focused on monitoring the brain activity of solitary individuals, Hyperscanning studies provide insights on inter-brain synchrony, advancing the field of social neuroscience.

Rodents are often used as a model to study social interactions due to their prosocial nature, tendency towards group living, and hierarchical behavior that has similarities to human behavior^12–15^. Over the years, several experimental paradigms have been developed to investigate behavior in physiological and pathological conditions, contributing to the understanding of social dynamics^16–21^. Consequently, rodents constitute an excellent animal model for developing experimental paradigms that deepen the understanding of behavioral mechanisms related to the social sphere.

The main difficulty in investigating the neuronal circuitry during the execution of social interaction repertoire is mainly linked to the need to keep the animals in conditions of free movement. This limit was overcome with the advent of miniaturized optical systems which made it possible to monitor brain activity up to cellular resolutions^22–30^. Although the progress of in vivo optical imaging techniques has made it possible to simultaneously monitor the neuronal activity of two mice involved in social behaviors^31,32^, whether there is a possible inter-brain cortex-wide synchrony in freely-moving mice involved in social behavior is still under debate.

In this study, we present an integrated platform for conducting Hyperscanning studies in awake freely moving mice. First, we developed a head-mounted Miniaturized Cerebral microScope, the “MiCe-μScope”, capable of monitoring neural activity from the dorsal portion of the mouse cortex with performance comparable to a regular tabletop microscope. Second, we designed an experimental task in which the animal behavior is modulated by social interaction. Third, through simultaneous recordings of mice engaged in prosocial behaviors, we show that social interaction modulates synchrony between the two brains of the interacting animals in two different frequency bands. Our work provides an experimental framework for conducting Hyperscanning studies in mice capitalizing on the genetic and invasive tools developed in this model to uncover the neural substrates responsible for social behavior.

## RESULTS

### The MiCe-μScope records wide-field calcium imaging in awake freely moving mice

To study the cortical activity of socially interacting animals, we developed a new device, the MiCe-μScope (Miniaturized Cerebral microscope for mice), which allows monitoring neural activity in awake and freely moving mice. This system provides the capability to observe and analyze neural processes across the entire cortical mantle.

The MiCe-μScope components are built using stereo-lithography 3D printing technology. It is equipped with two blue LEDs (with a dominant wavelength between 465 and 485 nm), paired with a circular excitation filter (ET 470/40X) that selects the correct wavelength to excite the GCaMP6f indicator. An emission filter (ET 525/50m) is then inserted before the camera to filter the fluorescence signal. Finally, a complementary metal-oxide-semiconductor imaging sensor (Spy camera for Raspberry Pi 5MP) is applied on top of the device to record cortical activity through the intact skull over the entire cortical mantle (Figure 1A) (see MiCe-μScope in Methods). The optical device is mounted on the mouse’s head using an implant ring with an aluminum post (head-post) and a set of screws for fixation (Figure 1B). This setup ensures perfect mechanical coupling between the MiCe-μScope and the animal’s head, allowing for manipulation of the animal and device mounting without the need for anesthesia. The system weighs less than 3 grams (2.9 grams), has a sample frequency of 25 Hz, and provides a wide field of view (FOV) of approximately 10.5 mm, sufficient to monitor both hemispheres simultaneously. The symmetrical illumination from the two blue LEDs ensures homogeneous lighting across the entire FOV. The average resolution along the FOV, calculated using the 1951 USAF resolution test target, is (31.00 ± 5.01) (Supplementary Figure 1A-B-C).

**Figure 1.**
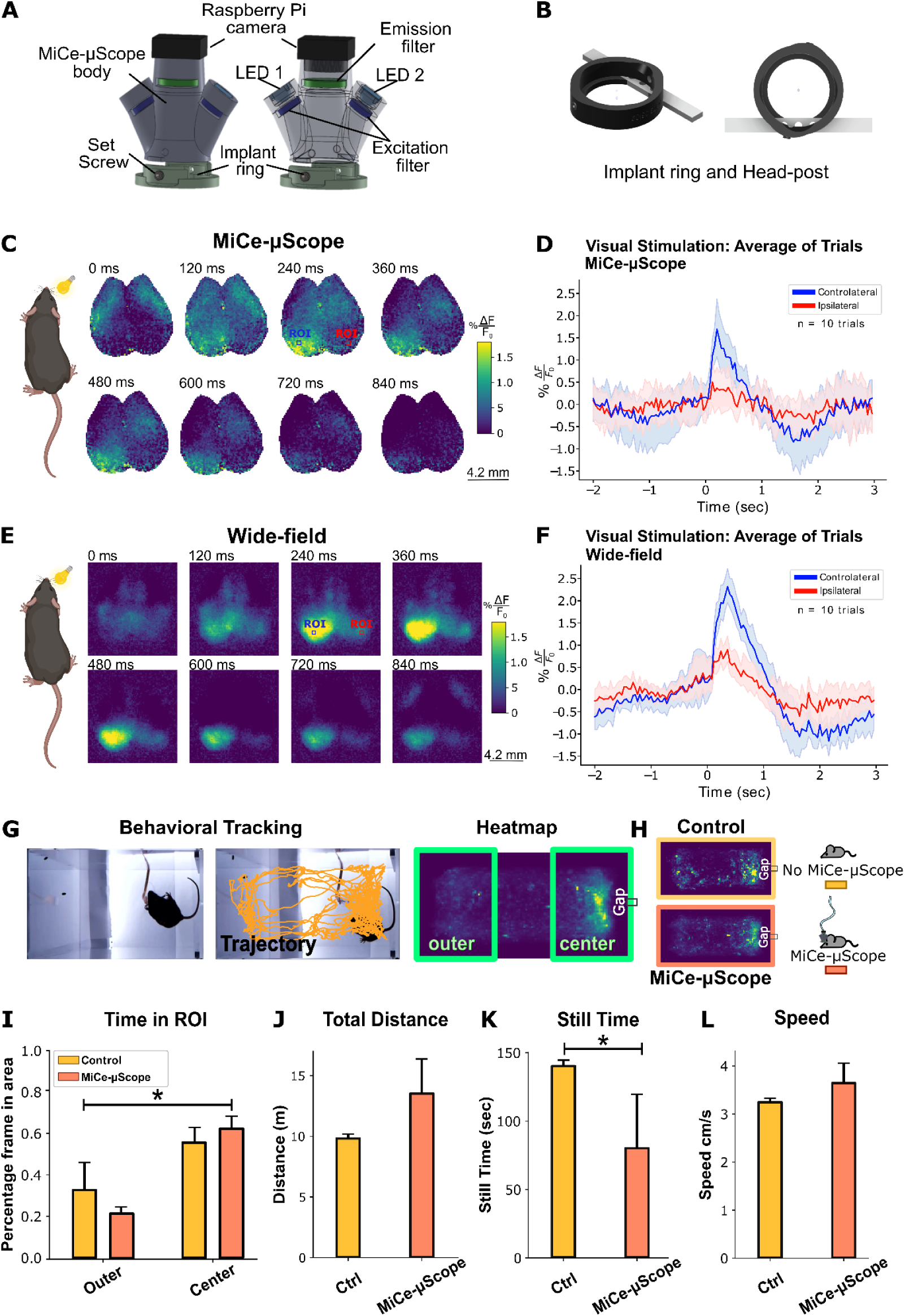
MiCe-μScope allows monitoring of cortical activity in awake freely moving animals. (A) MiCe-μScope: rendering of the MiCe-μScope with all optical and mechanical components. (B) Rendering of the head-post with the implant ring. (C-E) Cortical responses to visual stimulation: cortical responses evoked by visual stimulation recorded with the MiCe-μScope e wide-field microscope. Mean activations sequence distributed over both hemispheres following visual stimulation of the right eye using white light (duration of pulse = 100 ms, n = 10 trials in 1 mouse) (reported from the onset to 840 ms post-stimulus). Blue and red ROIs indicate respectively the contralateral and ipsilateral cortical regions of the visual cortex from which the fluorescence signals were extracted using the MiCe-μScope (C) and the conventional wide-field microscope (E) (Scale bar: 4.2 mm). (D-F) Average cortical activity: plot shows the average cortical activity (expressed as % ΔF/F_0_) of the 10 visual stimuli extracted from the visual cortex of the contralateral (blue) and Ipsilateral (red) hemisphere using both optical system: MiCe-μScope (D) (Contralateral hemisphere: 1.70 ± 0.53 %; Ipsilateral hemisphere: 0.51 ± 0.69 %) and the conventional wide-field microscope (F) (Contralateral hemisphere: 2.32 ± 0.55 %; Ipsilateral hemisphere: 0.90 ± 0.53 %). The highlighted line represents the mean, and the shadow represents the 95% confidence interval. (G) Behavioral box setup: bottom view representation of the behavioral box containing the mouse with the MiCe-μScope. The black mouse on a white background was used to apply a custom-made algorithm for detecting the animal’s centroid and performing animal tracking to generate the heatmap. (H) Heatmaps: heatmaps depict the time animals spent in different parts of the arena (Outer and Center) in two experimental conditions: with the MiCe-μScope and without Control. (I) Time in outer and center regions: plot shows the time animals spent in the two Outer and Center regions in the two experimental groups: those wearing the MiCe-μScope and the Control group (Control center: 0.61 ± 0.15; Control outer: 0.28 ± 0.13; MiCe-μScope center: 0.62 ± 0.05; MiCe-μScope outer: 0.22 ± 0.04; *p < 0.001, Mixed Effect ANOVA). (J) Total distance: plot shows the distance traveled by the animals with and without the MiCe-μScope inside the behavioral box (Control: 9.90 ± 0.27 m; MiCe-μScope: 13.14 ± 3.22 m). (K) Time in quiet state: plot represents the time animals spent in a quiet state in the two experimental groups: MiCe-μScope and Control (Control: 139.64 ± 5.00 sec; MiCe-μScope: 79.55 ± 51.76 sec; *p <0.05, Welch t-test). (L) Speed of animals: plot represents the speed of the animals with and without the MiCe-μScope in the behavioral box (Control: 3.23 ± 0.08 cm/s; MiCe-μScope: 3.62 ± 0.58 cm/s). The bar plot represents the average value, and the error bars represent the 95% confidence interval.

The miniaturized structure of the MiCe-μScope, as depicted in Figure 1A-B, enables a more natural and unrestricted observation of behavioral dynamics, contributing to a comprehensive exploration of the neural mechanism underlying social behaviors in rodents. First, we tested if the MiCe-μScope can successfully record neural activity in GCaMP6f transgenic mice. To this end, a visual stimulation experiment was performed, and compared the results of this experiment with those obtained using a conventional epifluorescence wide-field microscope (Figure 1C-D)^33,34^. Cortical activity was monitored on both hemispheres of a mouse while delivering a short pulse of light (100 ms) on the right eye (n = 10 trials in 1 mouse). Both the MiCe-μScope and the wide-field microscope detected a cortical response in the contralateral left visual cortex (Figure 1C-D) following the presentation of the visual stimulus. The maximum activation recorded after stimulus onset is comparable between the two optical systems: (228 ± 65) ms for MiCe-μScope and (384 ± 15) ms for the wide-field microscope (Supplementary Figure 1D and G). Notably, at the single-trial level, the evoked response can be observed mainly in the contralateral hemisphere and is very noisy in the ipsilateral hemisphere (Supplementary Figures 1E-F-H-I). When averaging together the responses to all stimuli, the amplitude of the evoked response (% ΔF/F) in the selected ROI was 1.70 ± 0.53 % for the MiCe-μScope (Figure 1D) and was again comparable to the values obtained with the conventional wide-field microscope, 2.32 ± 0.55 % (Figure 1F) in the contralateral hemisphere. In both optical systems, the response recorded by the visual stimulus in the ipsilateral hemisphere is lower than in the contralateral one (MiCe-μScope: 0.51 ± 0.69 %; wide-field: 0.90 ± 0.53 %; Figure 1F). Therefore, despite differences in sensor performance, optical components, and resolution between the two devices, the consistency in the observed cortical response and the comparable activation patterns between the MiCe-μScope and the conventional wide-field system indicate the successful validation and characterization of this novel tool. Therefore, the MiCe-μScope is a reliable tool for monitoring and measuring a stimulus-evoked response with qualitative and quantitative characteristics comparable to those of the wide-field microscope.

To test whether the system mounted on the rodent’s head could potentially influence its behavior, we compared the exploratory abilities of animals with (MiCe-μScope group) and without (Control group) the miniature microscope inside the behavioral box used for the social interaction test. Briefly, the behavioral box consisted of a rectangular plexiglass arena separated in the middle by a transparent barrier with a small aperture (gap, 1 by 10 cm) to facilitate mouse-to-mouse interaction in the center of the box (see Behavioral Setup in Methods). Videos of the behavioral recordings were manually analyzed to extract the trajectories and the heatmaps of the position of the animal (Figure 1G). The presence of the MiCe-μScope did not hinder the mice exploratory behavior (Figure 1I-J-K-L). No statistical significant differences were observed between Control and MiCe-μScope group when considering the time spent in the outer or center portion of the arena (Mixed Effect ANOVA, factor MICE-μSCOPE F(6,1) = 1.6, p = 0.252; Figure 1I) demonstrating that the presence of the device on the rodents’ head does not affect their ability to explore the behavioral arena. (Figure 1I). To further validate the device, we analyzed the distance traveled by the mice in both experimental groups. The results indicate that mice wearing the MiCe-μScope covered, on average, a greater distance compared to the control group (Control: 9.90 ± 0.27 m; MiCe-μScope: 13.14 ± 3.22 m; Figure 1J) although not statistically significant (Welch t-test, t(5) = 2.4, p-value = 0.057; Figure 1J). To evaluate if the weight of the MiCe-μScope could impose limits on the locomotion behavior, we computed the time in which the animals remained still. Unexpectedly, the control mice stayed still longer than those wearing the device (Control: 139.64 ± 5.00 sec; MiCe-μScope: 79.55 ± 51.76 sec; Figure 1K; Welch t-test, t(5) = 2.8, p-value = 0.036). Following the analysis of the animal’s quiet time, we shifted our focus to the period in which the animals were in motion. By calculating the average instantaneous speed per mouse expressed as an instantaneous velocity ± SD, we found that mice exhibited comparable instantaneous speeds (Control: 3.23 ± 0.08 cm/s; MiCe-μScope: 3.62 ± 0.58 cm/s; Figure 1L; Welch t-test, t(5) = 1.6, p-value = 0.156).

In addition, both groups of mice wearing the MiCe-μScope and not, tend to spend more time in the central region of the arena, particularly near the gap (Figure 1H). This preference for the central arena could be attributed to the structural asymmetry of the arena, which encourages animals to explore the gap and, as a result, spend more time in the center. The quantification of the area occupied by the animals in the two experimental groups, Control and MiCe-μScope, expressed as the average percentage of frame in the center (close to the gap) and outer regions (Control center: 0.61 ± 0.15; Control outer: 0.28 ± 0.13; MiCe-μScope center: 0.62 ± 0.05; MiCe-μScope outer: 0.22 ± 0.04; Figure 1I) confirms that both groups, with and without the MiCe-μScope, exhibit a significant preference for the central area over the outer region of the arena (Mixed Effect ANOVA, repeated factor REGION F(6,1) = 62.34, p<0.001; Figure 1I).

In conclusion, we have successfully developed a device with optical characteristics and mechanical stability that enables the monitoring of both cortical hemispheres through the intact skull in freely moving mice. The design and structure of the MiCe-μScope allow the device to be comfortably mounted on the heads of animals without the need for anesthesia and without hindering their motor and exploratory abilities. This demonstrates that the device has the appropriate characteristics and potential to investigate cortical dynamics simultaneously with behavior monitoring. Consequently, mice are free to move and exhibit their normal, natural behavior within the arena.

### Social interaction modulates the behavior of the animal in a modified Three-Chamber test

We developed a behavioral task to investigate both the neuronal and behavioral dynamics of social interactions. Taking inspiration from the behavioral paradigms of the Three-Chamber test^35^ and the linear chamber apparatus^36^, we designed a linear box (540 x 150 x 200 mm) with a central barrier equipped with a slot, referred to as the “gap” (10 x 10 x 100 mm) to facilitate the interaction between animals at the center of the apparatus. The social interaction test consisted of a 7-minute “interaction” phase, preceded by two “solitary” habituation phases of the same duration (Figure 2A). First, we analyzed how the animal’s behavior in the chamber is affected by the interaction, by comparing heatmaps of the mice position in solitary vs interacting conditions (Figure 2B-C). Mice tend to occupy the central region of the chamber independently of the presence or absence of interaction. However, in the Interaction condition the mice tend to get closer to the barrier, with a higher probability of them occupying the region in proximity to the gap compared to the outer region of the box (Figure 2D). Second, we examined the distance traveled by the animals in the two conditions. Solitary mice travel distances comparable to the interaction session (Solitary: 12.33 ± 3.11 m, Interaction: 11.52 ± 3.41 m; Mixed Effect ANOVA, factor CONDITION F(6,1) = 1.84, p = 0.224; Figure 2E). This suggests that even though all experimental groups tend to occupy the region near the gap more frequently when both animals are present, they tend to stay in this region, exploring the gap or interacting with their dyad partner.

**Figure 2.**
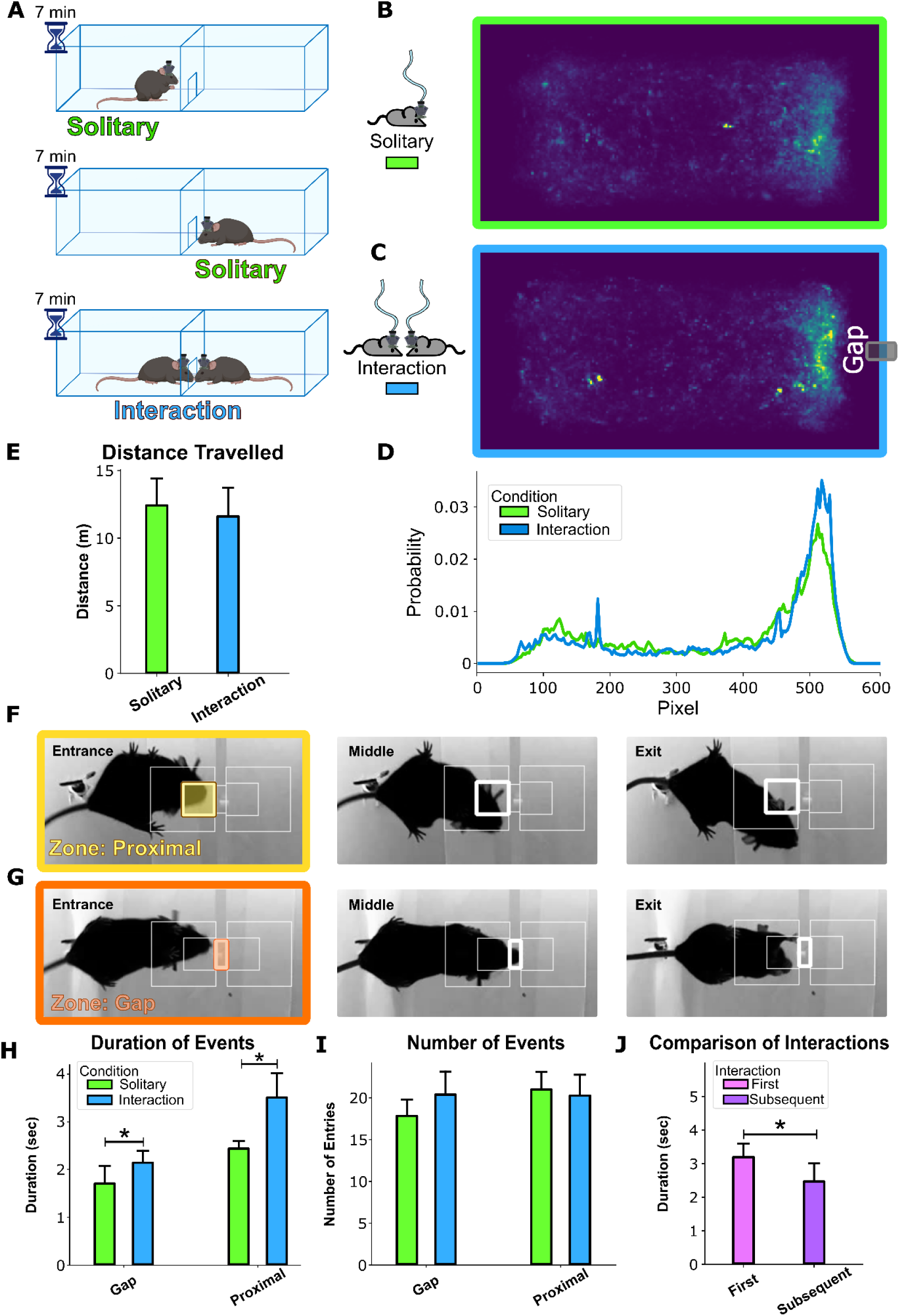
Social interaction influences the behavior of the animal in a modified three-chamber test. (A) Time sequence of the three experimental phases: the behavioral task includes a habituation session (Solitary condition) followed by an interaction session (interaction condition). Each session lasts 7 minutes. (B - C) Heatmaps of the behavioral box: heatmaps depict the time spent by mice in different areas during Solitary (B) and Interaction (C) conditions. (D) Probability of Occupancy Density: the graph shows the probability of occupancy density within the heatmaps in both Solitary and Interaction conditions. (E) Average distance traveled: the plot illustrates the average distance covered by mice during Solitary (12.33 ± 3.11 m; green) and Interaction (11.52 ± 3.41 m; blue) conditions. (F - G) Nose poke events: images represent nose poke events in Proximal (F) and Gap (G) zones, showing entrance, intermediate, and exit frames. (H) Duration of nose poke events: plot presents the duration of nose poke events in Gap and Proximal zones for Solitary (green) and Interaction (blue) conditions (Gap Solitary: 1.66 ± 0.45 s; Gap Interaction: 2.11 ± 0.26 s; Proximal Solitary: 2.41 ± 0.40 s; Proximal Interaction: 3.47 ± 0.45 s; * p < 0.001, Repeated Measures ANOVA). (I) Number of nose poke events: plot represents the average number of entries within the Proximal and Gap zones for Solitary (green) and Interaction (blue) conditions (Gap Solitary: 17.64 ± 5.21, Gap Interaction: 20.48 ± 5.83, Proximal Solitary: 20.82 ± 6.32, Proximal Interaction: 20.25 ± 4.73). (J) Comparison of interactions: comparison between the average duration of the first interaction and all subsequent interactions (Duration first interaction: 3.15 ± 0.59 s; Duration of subsequent interactions 2.42 ± 0.79 s; * p < 0.05, Paired t-test). All bar plots represent the average value while the error bar represents the 95% confidence interval.

Since the heatmaps suggest differences in animal behavior near the barrier, we turned our attention to the central region of the box. In particular, we defined two regions near the barrier: “Proximal” and “Gap” (Figure 2F-G). Mice spent more time in both the Gap and the Proximal zone when there was another animal in the opposite part of the arena (Repeated Measures ANOVA, factor CONDITION, F(7,1) = 33.36, p<0.001; Figure 2H). These findings highlight how the presence of another animal in the arena influences the amount of time spent in these specific zones during nose poke events. When quantifying the average number of entries in the Gap and Proximal zones in both the Solitary and Interaction conditions (Gap Solitary: 17.64 ± 5.21, Gap Interaction: 20.48 ± 5.83, Proximal Solitary: 20.82 ± 6.32, Proximal Interaction: 20.25 ± 4.73; Figure 2I), except for a slight increase in the number of entries to the Gap in the Interaction condition compared to the Solitary condition, no statistically significant differences were observed in the number of events between the zones and the different conditions (Repeated Measures ANOVA, factor CONDITION, F(7,1)=1.35, p = 0.283; Figure 2I). Thus, from the analysis of the number of entries, it becomes evident that all animals, regardless of whether or not a second mouse is present on the other side of the barrier, tend to exhibit a comparable number of nose poke events in both the Gap and Proximal zones. However, the presence of a second animal on the other side of the barrier significantly influences the duration of the nose poke events in these two identified zones (Gap and Proximal). This emphasized the significant effect of the presence of a dyadic partner on interaction events.

The experimental paradigm used in this study involved forming dyads in which each animal encountered a novel animal during each session, without any previous contact. To understand how the animal’s behavior changed across different interaction events, the time elapsed during the first interaction event and compared with subsequent ones (Figure 2J). In the dyadic sessions, mice performed an average (5.36 ± 3.05) interactions per session. Additionally, they spent more time at the gap during their first interaction (3.15 ± 0.59 s) compared to subsequent events (2.42 ± 0.79 s; Figure 2J). The difference between the duration of the first interaction and the subsequent ones is statistically significant (Paired t-test, t(7) = 3.29, p-value=0.013; Figure 2J). This suggests that there is a notable change in behavior between the initial interaction and the subsequent one within the dyad. This could indicate that the initial encounter between two unfamiliar mice at the gap was characterized by prolonged interactions, suggesting that the novelty and lack of previous contact in the social arena may have influenced their behavior.

In conclusion, we can affirm that although the structural asymmetry of the behavioral arena encourages both solitary and interacting mice to explore the zone near the barrier, it’s when both animals are present that they tend to spend more time near the gap compared to the solitary condition. Based on the results obtained from the behavioral analyses, we can validate the functionality of the arena as a behavioral test to investigate the dynamics of social interaction.

### Social interaction enhances inter-brain synchrony in the slow and infra-slow frequency bands

Since the behavioral results show that interactions occurred at the gap region of the behavioral box, we wondered if inter-brain coupling or synchronization occurred when mice were in this region. To this aim, inspired by Hyperscanning studies in humans^8,10,11^, we analyzed inter-brain coupling by computing the Wavelet Coherence Transform (WCT) of the two fluorescence signals extracted from each mouse of the dyad over the whole cortex. WCT is a method commonly used to investigate the coupling between two signals in both time and frequency domains, and it is particularly suitable for studying complex, dynamic interactions, such as social behaviors. The WCT was computed for the whole interaction session and segmented based on two different behaviors observed at the gap: Gap Interacting and Gap Non-Interacting. Importantly, each coherence spectrum estimated from the two behavioral conditions was also tested against surrogate data generated from the solitary behavioral sessions named Gap Surrogate (refer to the “Behavioral Analysis” section in Methods).

The wavelet analysis facilitated the generation of coherence spectra as a function of frequency. The plots in Figure 3A-B, depict the average coherence spectrum (n = 9 dyads) for the entire cortex for the Gap Interacting and Non-Interacting conditions versus the Gap Surrogate control condition. The full-spectrum highlights the existence of two frequency bands with increased coherence: the first in the infra-slow range (1/16 -¼ Hz), when comparing Gap Interacting and Gap Non-Interacting vs Gap Surrogate, and the second in the Slow range (2 - 4 Hz) only when comparing the Gap Interacting vs the Gap Surrogate. Therefore, considering only these two frequency ranges, for each dyad we extracted and compared the peak coherence in each of the three behavioral conditions. In this way, we were also able to allow each dyad to have a different frequency peak as opposed to averaging the coherence among the entire band. In addition, this approach enabled us to calculate the frequency of the maximum coherence peak within the spectrum. When considering the infra-slow band, we found that the peak coherence in this band was modulated by the behavioral condition (Repeated Measures ANOVA, factor BEHAVIOR, F(16,2) = 9.58, p = 0.002). Specifically, we found the highest value of the peak coherence during the Gap Interacting (0.47 ± 0.15) followed by the Gap Non-Interacting (0.36 ± 0.04), and, finally, by the Gap Surrogate (0.29 ± 0.05) with all comparisons among the three conditions being statistically significant (Paired t-test with Holm-Bonferroni correction, t(8) = 2.34, p=0.048; t(8) = 3.43, p=0.018; t(8)=4.20, p=0.009, Figure 3C). Similarly, In the slow band, the peak coherence was significantly modulated by the behavioral condition (Repeated Measures ANOVA, factor BEHAVIOR, F(16,2) = 12.99, p < 0.001). However, the peak coherence in the Gap Interacting condition (0.30 ± 0.04) was significantly greater than the one in both the Gap Non Interacting (0.25 ± 0.01) and the Gap Surrogate (0.24 ± 0.03) behaviors (Paired t-test with Holm-Bonferroni correction, t(8) = 3.57, p=0.015; t(8) = 4.29, p=0.008, Figure 3E). In contrast, there was no significant difference between the Gap Non-Interacting and the Gap Surrogate (Paired t-test with Holm-Bonferroni correction t(8) = 1.32, p=0.223, Figure 3E).

**Figure 3.**
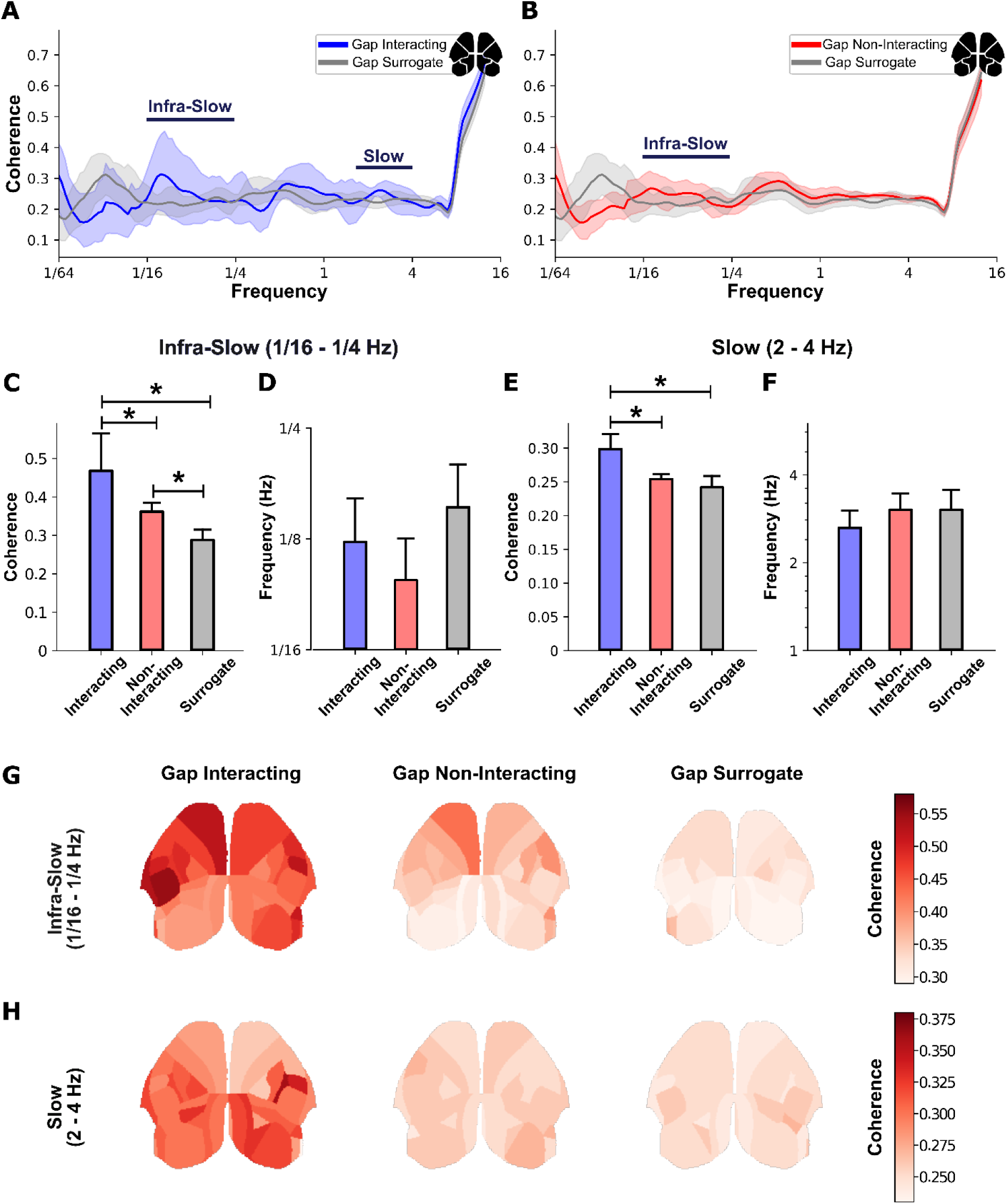
Social interaction increases slow and infra-slow frequency band inter-brain synchrony. (A) Full-spectrum coherence across the entire cortex during Gap Interacting and Gap Surrogate. Lines represent frequency bands with a statistically significant increase in coherence (1/16 - ¼ Hz, infra-slow; 2 - 4 Hz, slow) (Number of dyads = 9). (B) Full-spectrum coherence across the entire cortex during Gap Non-Interacting and Gap Surrogate (Number of dyads = 9). The highlighted line represents the mean, and the shadow represents the 95% confidence interval. (C) Graphs showing the average coherence of the maximum peaks of each dyad extracted in the infra-slow frequency band (1/16 - ¼ Hz) by behavioral event (Gap Interacting = 0.47 ± 0.15; Gap Non-Interacting = 0.36 ± 0.04; Gap Surrogate = 0.29 ± 0.05; n=9, * p < 0.05 Paired t-test with Holm-Bonferroni correction). (D) Graphs showing the average infra-slow frequencies where the maximum peaks of coherence are located by behavioral event (Gap Interacting = 0.12 ± 0.06; Gap Non-Interacting = 0.10 ± 0.04; Gap Surrogate = 0.15 ± 0.08; n=9). (E) Graphs showing the average coherence of the maximum peaks of each dyad extracted in the slow frequency band (2 - 4 Hz) by behavioral event (Gap Interacting = 0.30 ± 0.04; Gap Non-Interacting = 0.25 ± 0.01; Gap Surrogate = 0.24 ± 0.03; n=9, * p < 0.05 Paired t-test with Holm-Bonferroni correction). (F) Graphs showing the average slow frequencies where the maximum peaks of coherence are located by behavioral event (Gap Interacting = 2.63 ± 0.59; Gap Non-Interacting = 3.04 ± 0.68; Gap Surrogate = 3.03 ± 0.90; n=9). The bar plot represents the average value, and the error bar represents the 95% confidence interval. (G - H) Maps of the cortical mantle parceled according to the Allen Institute Mouse Brain Atlas showing the maximum peak of coherence in the two frequency bands (infra-slow and slow) by behavioral event (Gap Interacting, Gap Non-Interacting, and Gap Surrogate).

As stated earlier, we tested if the frequency of the coherence peak was modulated by the three different behavioral conditions for both the slow and the infra-slow frequency ranges. For both frequency bands, the behavior weakly modulated the value of the peak frequency leading to non-statistically significant changes (Repeated Measures ANOVA, factor BEHAVIOR, F(16,2) = 1.65, p = 0.223 and F(16,2) = 0.93, p = 0.416 for the infra-slow band and Slow band respectively; Figure 3D and F).

The coherence analysis on calcium imaging data suggests that neural activity is synchronized across the brains of interacting mice at least in two different frequency bands. This finding indicates that the neural activity in both brains is coupled and coordinated over the entire cortex. The level of coherence is strongly influenced by whether social interaction is occurring at the gap.

Following the observation of this distributed coherence variation over the entire cortex, we were interested in defining which cortical regions contribute to the synchronization induced by social behavior in the slow and infra-slow frequency bands. In this context, WCT analysis was performed by parceling the entire cortex based on Allen’s Mouse Brain Atlas (Supplementary Figure 2). Figure 3G-H shows the maps of the maximum coherence peaks on a colorimetric scale for all dyads in the three behavioral conditions (Gap Interacting, Gap Non-Interacting, and Gap Surrogate). The results from the maps confirm that, in both frequency bands, the maximum coherence peaks are observed in the Gap Interacting behavior condition, followed by the Gap Non-Interacting and Surrogate conditions (Figure 3G-H). However, the analysis of the maps reveals a differential involvement of individual cortical areas. In the Gap Interacting condition, the areas that contribute mostly to the coherence are the motor and somatosensory areas in the infra-slow band while in the slow band, a contribution emerges mainly from retrosplenial and visual areas. The involvement of the motor and somatosensory areas persists, albeit to a lesser extent, even within the infra-slow band during the Gap Non-Interacting condition. Consistently with the results obtained from the entire cortex, the analysis of individual areas in Surrogate dyads shows low coherence, confirming the absence of inter-brain coupling between the dyads. This reinforces the idea that the observed inter-brain coupling is linked to the social behavior exhibited by the interacting mice.

The analysis of coherence conducted on each cortical region suggests the presence of two different coherence maps for the infra-slow and the slow frequency bands that are modulated by social behavior.

## DISCUSSION

In this study, we developed a miniaturized optical system, the “MiCe-μScope”, to conduct wide-field calcium imaging in awake freely moving mice engaged in social behavior. To this end, we devised a behavioral paradigm that enabled the investigation of behavioral dynamics during social interactions. By combining the optical system with this behavioral paradigm, we were able to simultaneously explore cortical activity across both hemispheres in dyads of freely moving mice engaged in social interaction tasks. This approach revealed that social interactions modulate inter-brain synchrony between the two interacting mice across different frequency bands and that the synchrony engages specific cortical networks upon each frequency band.

### Comparisons with other miniaturized microscopes

The MiCe-μScope possesses the mechanical and optical characteristics necessary for monitoring neuronal activity across the entire cortex with good stability. Its functionality was confirmed through visual stimulation experiments, where responses evoked by the stimulus were compared with those recorded by a conventional wide-field microscope. While variations exist in sensor performance and optical resolution between the two systems, the clear and distinguishable response in calcium activity at the single trial level underscores the MiCe-μScope’s efficacy in recording fluorescence changes induced by sensory stimuli. This validation ensures that the MiCe-μScope reliably captures distributed cortical activity, akin to wide-field systems in experiments conducted under head-fixed conditions^26,34,37–41^.

Over the past two decades a significant push in generating design miniaturized optical devices that allow monitoring neuron activity in awake freely moving animals to investigate neural processing in underlying ethologically significant behaviors^22–26^. These systems enable real-time monitoring of neural activity during natural and unrestrained behaviors such as social exploration, social interactions, and locomotion^22,42,43^. The great majority of the existing systems are designed to achieve cellular resolution imaging of small brain regions by creating a cranial window and using a GRIN lens at the target site^22–25,31^. Our Miniscope, along with the one developed by Rynes and colleagues^26^, stands out as the only device capable of wide-field calcium imaging in awake freely moving animals. Although these two Miniscope offer a large field of view, good optical resolution for capturing wide-field calcium imaging, and both generate imaging data comparable to those obtained from conventional large tabletop fluorescence microscopes, several important differences exist and must be taken into account between the two devices. Unlike the system developed by Rynes and colleagues^26^, our system does not include a dual stroboscopic illumination system for performing hemodynamic correction of the fluorescence signal. However, this omission allowed us to develop a miniaturized optical system weighing less than 3 grams, which except for the MiCe-μScope body and the implant ring, is composed of components readily available over the shelf and simple to integrate. Another key difference is that the MiCe-μScope performs imaging through the intact skull while the one designed by Rynes and colleagues is mainly built to use a transparent See-Shells polymer directly in contact with the dorsal surface of the cerebral cortex. In this way, while the transparent polymer allows for high-resolution functional and structural imaging^44^ we decided to avoid the need for invasive procedures such as thinning the skull or performing craniotomies, thereby reducing the complexity of surgical procedures, the inflammatory responses typically associated with surgical procedures, and minimizing recovery times for the animals. Finally, the lightweight implant ring with its integrated aluminum head-post simplified animal handling, allowing rapid attachment of the MiCe-μScope in seconds without the need to anesthetize the animal.

The MiCe-μScope’s lightweight design, together with the result that this device did not alter the mice’s behavior, as shown in Figure 1, makes it an ideal tool for studying neural activity in awake freely moving animals. This information is crucial to ensuring that MiCe-μScope recordings accurately represent the natural behavior of animals during social interaction experiments.

### Social interaction behavioral paradigm

A widely employed behavioral paradigm for studying social behavior in mice is the three-chamber apparatus, commonly used to investigate behavioral approaches^45–49^. Recently, the linear chamber test was introduced as a variant for studying social approach behavior in mice^36^. Inspired by this variant, we developed a new version of the behavioral apparatus to investigate social interactions, with a specific focus on the social approach involving nose contact. Our behavioral apparatus included a transparent barrier with a central gap to facilitate social contact with the nose. In contrast to the three-chamber apparatus, which features two lateral chambers, our design aligns with the approach of the linear chamber test, aiming to reduce environmental exploration mechanisms and maximize the number of nose-pokes at the gap^36^. Unlike studies where social targets were confined to limited spaces^21,35,36,50^, our apparatus, similar to the study of Kingsbury and colleagues^31^, allowed both animals of the dyad to move freely within the behavioral setup without forcing the interaction between the two animals. The combination of the MiCe-μScope with our behavioral setup enabled us to simultaneously monitor the activity of the entire cortical mantle throughout the behavioral repertoire.

In the three-chamber test, the duration of time spent in the two side chambers is the most important and widely accepted parameter to assess sociability and preference for social novelty in rodents^21,35,50,51^. Similarly, in our behavioral task, social interaction modulated the time spent by an animal at the gap in the presence of a partner. Moreover, data showed no significant difference in the number of gap entries between solitary and interacting conditions which is in line with what has been observed in the three-chamber test for the number of entries^50^. Together, these results align with findings from established paradigms in the study of social approach^35,36,50^, thereby validating the efficacy of our behavioral apparatus as a model for investigating social interaction.

Despite the three-chamber test demonstrating comparable sociability results between male and female mice^50^, the choice to use female non-sibling dyads in our peer-social interaction study was influenced by previous findings suggesting that female mice, particularly of the C57BL/6J strain, exhibit a high level of sniffing and social exploration^52^. This decision aimed to foster gap-sniffing events and minimize the likelihood of aggressive behavior. Considering the possibility of conducting a similar study with male mice in dyads suggests the potential for exploring sex-related behavioral differences in the context of peer-social interaction.

### Synchrony among brains of interacting mice

By combining the MiCe-μScope with an optimized behavioral paradigm, we simultaneously investigated neuronal activity distributed across the entire cortex and the behavior of awake, freely moving mice dyads engaged in social interaction. The ability to monitor cortical activity in two animals simultaneously allowed the extraction and analysis of images showing fluorescence changes related to calcium activity, enabling the evaluation of the degree of inter-cerebral coupling between interacting mice. While similar practices are commonly conducted in humans in Hyperscanning studies^6,8,10,53,54^, social interaction experiments in animals are usually conducted recording one area in one of the interacting animal^2,3,36^ or, in minor part, recording small homotopic areas in both animals^31,55^ or large portion of the cortex in head restrained animals^32^. This work is the only one that employs wide-field calcium imaging over the entire dorsal portion of the cortex in couples of awake freely moving socially interacting animals. The extension of this methodology to animal models opens up new avenues for translational research. Utilizing these translational animal models could investigate how inter-brain synchrony in many cortical regions is modulated in conditions where social interactions are impaired, such as autism spectrum disorder (ASD), depression, or social isolation.

In both humans and animals, the hallmark of social interaction is the synchronization of different brain regions^30,31,54,56–58^. The presence of inter-brain coupling between interacting subjects indicates that their neural activity is synchronized and coordinated during social behavior. This synchronization underscores the complexity of communication and coordination at the neural level during social interactions. Kingsbury and colleagues clearly show in mice the existence of inter-cerebral correlation of neurons in the prefrontal cortex, which varies depending on the nature of social interaction^31^. In addition, they reveal that the neural activity of one animal during interaction can be used to predict the behavior of its interacting peer. These findings suggest that synchronization is the expression of the mutual prediction performed by neurons of the medial prefrontal cortex (mPFC) in animals of interacting dyads. This highlights the role of the mPFC in mediating social interactions and coordinating neural activity between interacting individuals. Our study sought to build on this information, aiming to investigate neuronal dynamics across the entire cortex to evaluate the potential existence of inter-cerebral synchrony in all cortical regions in interacting mouse dyads. Our results show the existence of inter-brain synchrony involving frontal regions but extending over the entire cortical mantle. Therefore this synchrony could be the expression that the mutual prediction model is not limited exclusively to the mPFC but that there is a broader involvement of the entire cerebral cortex. Indeed, neuronal activation across the entire cortical mantle is not unique to social interaction but is a phenomenon observed in various behavioral tasks. Studies employing wide-field microscopy techniques have shown that neuronal activity associated with behavioral tasks leads to global activation of the cortex rather than being confined to specific anatomical regions^34,37,38,59–61^. This widespread activation reflecting the distributed nature of information processing performed by cortical regions can be therefore extended to social interaction. The involvement of other cortical areas in social interaction may be coordinated with the activation and information processing carried out by the mPFC, a region strongly implicated in social behavior^36,62–64^.

The synchronization of neural activity across different frequency bands during social interactions has been widely documented in the literature. This synchronization spans a wide range of frequencies, from theta to high-gamma oscillations^54,58,65–69^, highlighting the complexity of neural dynamics involved in social behavior. In our experiments, the relatively slow intrinsic dynamics of the calcium indicator^70^, as confirmed by simultaneous EEG recordings and wide-field calcium imaging^71^, limit our ability to directly assess whether inter-brain synchrony is present in the neural activity between two interacting mice in high-frequency bands. Nonetheless, the synchrony in the low-frequency bands observed here is important because the intrinsic communication bottleneck of the interaction implies that changes in synchronization in high-frequency bands are observed over slower time scales up to minutes^1^, highlighting the multiscale resolution of the phenomenon. Hence, the synchronization in the two frequency bands observed here could be the expression of information transfer and processing, between brains of interacting mice, that happens at two different timescales. In particular, the synchronization in the infra-slow frequency band could be at least partially associated with the movements of the whiskers or the paws^32^, while the synchronization in the low-frequency band, around 3Hz, could be an expression of respiratory-related neuronal activation behavior. It has been previously shown that respiratory rate is coupled with wide-spread cortical activity and induces oscillation in the delta range^72^ while synchronized 5Hz stimulation of awake freely moving mice increases sniffing-related social interaction behavior^73^.

The results reveal two distinct coherence maps corresponding to the infra-slow and slow frequency bands. The inter-brain synchrony observed in the infra-slow band likely reflects the engagement of both the motor and somatosensory systems, possibly due to coordinated movements of the upper limb, whiskers, and nose during social interactions. Indeed, a significant portion of prosocial behaviors, such as allogrooming, sniffing, and nose-to-nose contact, involve motor and somatosensory actions performed by interacting dyads^13,31^. In contrast, the inter-brain synchrony observed in the slow frequency band appears to be associated with the engagement of somatosensory areas and the associative components of visual and retrosplenial regions. This suggests the involvement of higher-order cognitive processes in social behavior, such as visual processing and spatial cognition. This could be compatible with mutual prediction models that occur in social interactions^1,31^.

In conclusion, the methods and the results shown in this work provide an experimental framework to conduct Hyperscanning studies over large portions of the cortex in an animal model that mirrors findings from human studies. By leveraging the genetic and experimental manipulations available in mice, future studies can fully exploit this framework to dissect the neuronal mechanisms underlying social behavior.

## METHODS

### Transgenic animals

All animal experiments conducted in this study were authorized by the Italian Ministry of Health (authorization 242-2020-PR). A total of 10 non-sibling female GCaMP6f mice (C57BL/6J-Tg(Thy1-GCaMP6f)GP5.17Dkim/J), were used to investigate peer social interactions with a novel animal, perform visual stimulation experiments and characterize the MiCe-μScope. To investigate social interactions, female non-sibling mice were specifically chosen. This decision was made to incentivize interactions between the mice while minimizing the likelihood of aggressive behaviors, which can arise more commonly in male mice. Mice were housed in a controlled environment with a 12-hour light/dark cycle with *ad libitum* access to water and food.

### Surgery

To visualize neural dynamics in vivo, mice involved in the experiments underwent surgery to create optical access and expose cortical regions of interest. The animals were anesthetized using inhalation of isoflurane 5% and were positioned in the stereotaxic apparatus (Model 1900, Kopf Instruments, Tujunga, California, USA) to ensure stability and precision during the surgical procedure. To maintain anesthesia throughout the surgery, the isoflurane concentration was adjusted to a range of 1-2%. Suppression of the nociceptive reflex is evaluated to determine the optimal depth of general anesthesia. To reduce the risk of hypothermia, mice were kept warm during the entire surgical procedure and until they fully recovered. A heating pad was placed under the ventral area of the mouse to maintain a stable body temperature. Continuous monitoring of the body temperature was carried out using a rectal probe to maintain it within the range of 36.5-37°C (ThermoStar, RWD, San Diego, California, USA), ensuring physiological well-being of the animals and minimizing any adverse effects associated with hypothermia. During the surgical procedure, the respiratory rate of the mice was monitored by observing the chest movements of the mice to ensure that the respiratory rate remained within the optimal range of ∼55-65 respirations per minute. To prevent ocular dehydration and potential damage to the eye, an ophthalmic gel (Lacrilube 42.5% Liquid Paraffin + 53.7% White Vaseline, AbbVie S.r.l., Latina, Italy) was applied to the eyeball to create a protective gel film. The procedure continued with the removal of hair from the dorsal surface of the animal’s head to expose the surgical area and minimize the risk of infectious complications. The depilated area was disinfected using ethanol 70% and Betadine 10% solution (Betadine 10%, Meda Pharma S.p.A, Milan, Italy) three times to ensure a high level of sterilization and minimize the risk of contamination. To reduce phlogosis processes, a subcutaneous injection of corticosteroid anti-inflammatory (Dexamethasone sodium phosphate, 5mg/Kg) was administered. For local anesthesia, a solution of lidocaine chlorohydrate 2% was applied topically through subcutaneous inoculation to minimize discomfort and pain during the procedure. A flap of skin was carefully lifted using tweezers and a cut was made with scissors to create a window on the skull. This window, approximately 2-2.5 cm^2^, provided access to both hemispheres of the brain. To maintain the exposed surface hydrated, a solution of sodium chloride 0.9% was applied. Any remaining hairs were removed from the surgical area, and the surface was thoroughly dried. Using the stereotaxic apparatus, two anatomical landmarks were identified and stained: Bregma (the midline bony landmark where the coronal and sagittal) and Lambda (the junction point between the two parietal bones and the occipital bone), which are generally 4.2 apart.

A layer of transparent dental cement (Super-Bond Universal Starter Kit, Sun Medical, Furutaka, Japan) was applied over the entire cortex. Great care was taken to ensure that the edges were meticulously covered, and no tissue flap was left exposed to the air to prevent the risk of infection. Subsequently, the implant ring with the head-post was placed on the animal’s head. To determine the correct position of the implanted ring, two critical anatomical features were considered. The first feature was the inclusion of the inferior cerebral vein on the rostral-caudal axis, and the second feature was the inclusion of the transverse sinus on the mid-lateral axis.

Another important parameter to check during the surgery was the parallel alignment of the implanted ring to the surface of the skull. Ensuring that the implanted ring was paralleled helped avoid introducing distortions in both the rostral-caudal and lateral axes. To secure the implant ring and the head-post, additional transparent dental cement was applied. This cement not only provided stability but also sealed the lateral margin between the ring and the skin to prevent any external light from entering the field of view during the imaging sessions. This step was essential to ensure high-quality imaging results without any interference.

At the end of the surgery, subcutaneous injection of analgesics and non-steroidal anti-inflammatory drugs (Carprofen, 5 mg/Kg) and 0.2 ml lactated Ringer’s solution (Lactated Ringers, Eurospital S.p.A., Trieste, Italy) was administered to support pain and hydration during the recovery period. During the postoperative hours, the mouse was monitored to assess the resumption of motor activity and responsiveness to stimuli.

To manage post-operative pain effectively, multimodal analgesic therapy was provided daily for the first three days following surgery. This therapy included subcutaneous administration of Carprofen (5 mg/kg) and Tramadol (20 mg/kg). Throughout the recovery period, which lasted one week, the mice were allowed free access to water and food to support their well-being and ensure a smooth recuperation from the surgical procedure.

### MiCe-μScope assembly

The MiCe-μScope, a miniaturized optical system developed within our laboratory, features a compact and lightweight design (22 x 20 x 12 mm, 2.7 grams). It consists of two main components: the body, which houses the sensor and optical elements, and an implant ring equipped with an aluminum head-post. The implant ring is affixed to the rodent’s head, simplifying the attachment of the MiCe-μScope body. The body and the implant ring were designed using computer-aided design (CAD) software called Autodesk Fusion 360. These designs were then produced using a stereo-lithography 3D printer (XFAB 2500HD; DWS System, Vicenza, Italy). The printer utilized a black photosensitive resin (Invicta DL385; DWS System, Vicenza, Italy) to create components that were isolated from external light sources, ensuring optimal imaging conditions.

The MiCe-μScope was equipped with two blue LEDs (CREE XPEBBL-L1-0000-00301; Cree, Inc., Durham, North Carolina, USA) with a dominant wavelength ranging between 465 and 485 nm. The two LEDs of the MiCe-μScope were connected in parallel using 36-gauge wires (Miniature Insulated Wire 36 AWG, UAA3601, Micron Meters; Tucker, Georgia, USA). Additionally, two resistors (10 Ohms ± 0.05% 0.25W, 1206; Stackpole Electronics, Inc., Raleigh, North Carolina, USA) were integrated into the circuit to ensure a balanced current supply. The entire illumination system was linked to a power supply capable of delivering 300 mA of current to the LEDs. These LEDs were paired with a circular excitation filter (450-490 nm, 4mm DIA, ET 470/40x; Chroma Technology Corp, Bellow Falls, Vermont, USA). The combination of the blue LEDs and the excitation filter facilitated fluorescence excitation, allowing for the visualization of neural activity in the optical window. The chosen wavelength range (between 465 and 485 nm) was essential to effectively excite the GCaMP6f indicator expressed in the excitatory cortical neurons. To filter the fluorescence signal and capture the emitted light, an emission filter (500-550 nm, 7mm DIA, ET 525/50m; Chroma Technology Corp, Bellow Falls, Vermont, USA) was inserted before the camera. This filter efficiently captured the fluorescence signal emitted by the excited neurons, while blocking unwanted light and background noise. A complementary metal-oxide-semiconductor (CMOS) imaging sensor (Spy Camera for Raspberry Pi 5MP, ID 1937 Adafruit Industries LLC, New York, New York, USA) was mounted on top of the MiCe-μScope device. This sensor allowed for the recording of cortical activity from both hemispheres simultaneously. The CMOS sensor of the MiCe-μScope was linked to a slip ring (Slip Ring mini 12 mm Dia, 12 wire, Adafruit Industries LLC, New York, New York, USA) located in the top panel of the behavioral setup. This system enabled the animals to move freely and rotate without any restrictions.

The combination of specific wavelengths, filters, and sensors enabled the MiCe-μScope to perform mesoscale imaging and record neural dynamics *in vivo*, at a sampling rate of 25 Hz, of the entire cortex through the intact skull.

### Behavioral setup

A modified version of the behavioral three-chamber apparatus was constructed to collect data on social interaction. A custom-made transparent plexiglass behavioral box (dimensions: 540 x 150 x 200 mm) was created, and inside it, a transparent barrier equipped with a gap in the middle (dimensions: 10 x 100 mm) was installed to divide the box into two identical compartments. The size of the gap allowed mice to interact with each other using their noses, while the transparent and open structure at the top ensured the maintenance of visual and olfactory contact with the partner. To capture behavior data at a rate of 40 Hz, a 5MP camera (5MP Camera, Okdo Design the World, London, United Kingdom) was placed beneath the behavioral box. For an optical field of view, the box was raised (approximately 60 cm) to ensure the edges were visible. The entire behavioral setup was enclosed by five custom-made white hardboard panels (two 900 x 750 mm for the front and back, two 750 x 450 mm for the sides, and one 900 x 450 mm for the top). To reduce sound interference during experiments, five sound-absorbing panels made of 50 mm thick polyurethane foam with pyramid-shaped patterns (Akustic Stop 100 x 100 x 5 cm 35 Kg/m2, NDA The Panels of Silence, Viterbo, Italy) were affixed to the external surface of the structure. To illuminate the entire experimental setup with white light, an LED strip (Kit Strip LED RGB 21, 6W 3m 24V, TMR Elcart, Milan, Italy) was applied along the internal perimeter of the ceiling. To prevent the light from disturbing the animals, paper sheets were applied to diffuse the light and create uniform illumination.

The behavioral camera and the slip rings linked with the CMOS sensor were connected via a flat cable (Cable FFC 15POS 1.00 mm 12”, Molex, Lisle, Illinois, USA) to the Raspberry Pi (Raspberry Pi4 Model B, 4GB; RaspberryPi, Caldecote, United Kingdom), and the videos of the animal’s movements and fluorescence calcium signals from the entire cortex were visualized on a Raspberry Pi Display (Display for Raspberry Pi LCD 7-inch touchscreen; RaspberryPi, Caldecote, United Kingdom). Finally, coaxial cables were used to synchronize all recording systems using a TTL signal generated by the data acquisition (DAQ) of a National Instruments.

#### Behavioral task

The experimental protocol developed for this study aimed to investigate the neural mechanism that coordinates social interactions in freely moving mice. After a recovery period following surgery to create an optical window on the dorsal surface of the skull, the animals were individually housed in an enriched environment within their home cages. This isolation was implemented to prevent potential aggressive behavior and injuries that could impact the experiment’s outcomes, particularly in the region with optical access to the cerebral cortex, which could impact the experiment’s outcomes. To encourage social interaction events, the mice were temporarily separated and placed in two distinct animal facilities for a minimum of two days before the commencement of the experimental sessions. This brief separation from potential social partners in separate facilities was intended to promote and stimulate social interaction events. The individual housing arrangements allowed for continuous visual and olfactory contact with conspecific in adjacent cages. Before the initiation of the recording sessions, the experimental conditions were established, and the interacting dyads were defined. Social interaction experiments were carried out twice a week, with a minimum of a two-day gap between each session. For these interactions, the dyads (animal A vs. animal B) were paired in such a way that during each recording session, animal A encountered a new animal B, and care was taken not to combine animals from the same litter. To ensure the best possible conditions, mice of each interacting dyad were individually housed in separate animal facilities for a minimum of two days before the experimental session. On the days of recording, mice of the interacting dyad were individually transferred from the animal facility to the experimental room and positioned on opposite sides of the room to minimize interaction before the session. The MiCe-μScope was then attached to the animals’ heads, after which they were placed in their cage for a habituation period of at least 15 minutes. This was done to facilitate the mice’s habituation to the presence of the device on their heads, ensuring their comfort and familiarity with the system before commencing the experimental task. Once the acclimation period was completed, the dyad components were introduced to the behavioral box to perform the social interaction task. The recording session was divided into three phases: in the first phase, animal A was placed alone in the behavioral box for 7 minutes. This allowed the animal to explore the environment and become accustomed to the experimental setup. It also aimed to reduce the exploration time during the interaction sessions. In the second phase, the same procedure was repeated for animal B. Finally, both animals (A and B) were placed in the behavioral box together to record the social interaction session for 7 minutes.

Throughout the entire experiment, cortical activity across the entire cortex and the behavioral responses of the animals were recorded concurrently using the MiCe-μScope and a Raspberry Pi camera positioned beneath the behavioral box. To transfer the animals from their home cage to the behavioral box, plastic containers were used. The animals were gently introduced into the containers, and upon placement in the behavioral box, they were allowed to move freely. During social interaction sessions, both animals were released into the behavioral box simultaneously. After each session, the behavioral box was cleaned with 70% ethanol, and the enclosure door was left open for a few minutes to allow the ethanol to evaporate and eliminate any residual animal scent from the recording session. Additionally, the plastic containers and all surfaces that came into contact with the animals were cleaned with 70% ethanol between each dyad to prevent olfactory contamination. Upon completing the recordings, the animals were returned to their respective home cages, housed in the animal facilities, and organized based on their dyads for subsequent sessions.

Thanks to the behavioral setup and the MiCe-μScope it was possible to simultaneously monitor both the behavior of the mice and the cortical activity distributed on both hemispheres in freely moving mice engaged in pair social interaction.

### Behavioral analysis

#### Behavioral tracking

The behavioral recordings were collected in *“.h264”* file format and subsequently converted into video files with the *“.avi”* extension. With the use of the ImageJ software, two ROIs were created to isolate the region of interest within the behavioral box. These ROIs are used as a mask to be applied to both the left and right compartments of the arena (size 300 x 150 mm, 600 x 300 pixels). These masks encompassed not only the central barrier containing the gap but also the edges of the behavioral box. Videos of the region of interest were employed for automated tracking of the animal’s positions using a custom-made blob detection algorithm. To ensure precise location tracking, maintaining a distinct contrast between the animal’s color and the background was crucial. To achieve this, aside from setting up an environment with white hard panels on the sides inside the behavioral setup, which enhanced the contrast between the black animal and the white background, a threshold was applied to the video files. This thresholding process transformed the RGB images into a binary video stack. The algorithm generated a circular blob around the animal, enabling the calculation of its centroid. This information was then used to determine the x and y coordinates of the animal‘s position within the box. To minimize errors in the algorithm’s evaluations and to encompass the animal’s entire behavioral repertoire, which might include activities like rearing or maximum stretching, two criteria were established: minimum and maximum area (500 and 50000 pixels respectively). These criteria served as limiting parameters for the tracking process.

The x and y coordinates obtained from the tracking algorithm were utilized to generate heatmaps, which are 2D histograms illustrating, on a colorimetric scale, the occupancy density of the animals in different groups and experimental conditions (bearing the MiCe-μScope, Control, Solitary, and Interaction condition). Pixel binning (200, 100) was also applied to the generated heatmaps, reducing the image resolution by ⅓.

The information derived from the tracking process enabled the quantification of various behavioral parameters for the animals. Notable parameters analyzed included the total distance traveled, calculated as the Euclidean distance, and the still time identified as the moment in which the animal’s speed was lower than 2.4 cm/s corresponding to a displacement of 0.06 cm per frame (25 ms). Finally, speed was calculated as distance traveled over time.

To assess the impact of the MiCe-μScope on the animals in the two experimental groups (bearing MiCe-μScope and Control), two additional regions were generated within the right and left compartments. These were defined as the Center region, with dimensions of 100 x 150 mm (200 x 300 pixels), located near the gap and extending for the entire length of the barrier, and the Outer region, of the same size but placed on the opposite side of the behavioral box compared to the gap. Combining the information obtained from the heatmaps and the internal subdivision of the behavioral box allowed for the quantification of the time spent in the two regions of the arena.

#### Gap analysis

To analyze the behavior of the mice in the Central region of the behavioral box near the barrier, a custom-made algorithm was employed to quantify events near the gap. The algorithm considered five rectangular regions, termed Zones, corresponding to the gap and near the barrier. Specifically, an area was defined in correspondence with the gap (Gap Zone) with dimensions (34 by 14 px). Two zones, situated on the right and left sides of the barrier and adjacent to the gap, were designed as Proximal Zones, with dimensions of (40 by 40 px). Finally, two additional zones were defined, encompassing the proximal region (80 by 80 px). Binary image sequences were utilized to quantify events, leveraging their high contrast between the background and the animal. The algorithm identified the presence of the mouse and the entry of the animal into the areas whenever (75%) of the pixels in the outlined regions transitioned from white to black. The analysis primarily focused on the identification of behavioral events, specifically nose poke events, in the Gap and Proximal Zones. To prevent false positives, such as entering areas with a tail or other behavioral phenomena, specific conditions were established to identify the events. To define entry into the Gap, the animal had to cross the Proximal Zone in preceding frames before entering the Gap Zone. Similarly, to define the entry into the Proximal Zone, the animal had to traverse the larger Zone that included the proximal region. Finally, all events were inspected visually by a human operator and eventual false positives were discarded.

The information derived from the analysis allowed us to identify both the duration and the number of nose poke events, categorized by zone in different experimental conditions (Solitary and Interaction). In the quantification, all events at the Gap that lasted less than 0.25 seconds (10 frames) were excluded.

#### Behavioral segmentation

Following the quantification of the animal’s behavior in the central region of the behavioral box, we proceeded to conduct a detailed analysis of their behavior in the area near the gap. Leveraging the behavioral quantification obtained from the algorithm, we utilized the information and events to construct an ethogram specific to this behavioral test. For this purpose, we employed BORIS, an easy-to-use event software designed for video coding and observation of various behaviors^74^. Specifically, we relied on systematic observation to investigate the behavior of the animals’ heads in the Gap zone during the three experimental sessions of the task.

The behaviors observed during the social interaction task were categorized into two main groups: Interacting Dyads (for the Interaction session) and Surrogate Dyads (for the Solitary sessions). Within the Interacting Dyads category, a further subdivision was conducted to distinguish between two different types of interaction. The first subcategory defined is Gap Interacting. Through a meticulous frame-by-frame examination of the video, the moments when both mice of the interacting dyad engaged in social interaction at the gap with their noses were identified. The onset of the event was determined as the first frame in which both animals initiated the social interaction event at the gap by entering the region near the gap with their nose. The end of the event was defined as the third frame in which one of the two animals exited the nose gap area. The second subcategory within Interacting Dyads is the Gap Non-Interacting condition. This behavior occurs when both animals are present in the behavioral box, but only one of the two members of the dyads approaches the gap to interact with their noses. In this case, the onset of the event was defined as the first frame in which the mouse enters the gap with its nose, and the end is defined as the third frame after the animal exits the gap. During this event, the other partner of the dyad could be in any region of the behavioral box.

Due to the intricate nature of the behavioral repertoire displayed by the animals within the behavioral box, categorizing these two types of behavior individually was not always feasible, as events were sometimes interconnected. Specifically, there were instances during the Gap Non-Interacting condition, where a single mouse had its nose inside the gap, and the animal on the other side of the barrier entered the region with its nose, initiating a behavioral event of social interaction, “Gap Interacting”. This could be triggered by a sudden approach of the animal to the barrier or because it was already very close to the gap. This occurred without the nose exiting the gap from the animals who had initiated the Gap Non-Interacting event. In such a case we promptly defined the end of the Gap Non-Interacting behavioral event (three frames before the start of the interaction) and initiated the Gap Interacting event on the next frame. This dynamic also applies to situations where, during a Gap Interacting event, one of the two animals decided to interrupt the interaction by moving its nose away from the gap, while the other remained with its nose in the gap and continued to explore the gap in solitude. This led to the initiation of a Gap Non-Interacting event. In this scenario, the end of the Gap Interacting event was defined three frames after the animal ceased the interaction by moving away from the gap, and in the next frame, we commenced the Gap Non-Interacting condition for the single animal who remained in the gap with the nose.

To conduct a comparative analysis of experimental results related to behavior segmentation, we aimed to establish a control condition that mirrored the specific behaviors observed in the interacting dyads. Thus, we decided to create Surrogate Dyads. This involved utilizing two single habituation sessions, during which the animal explored the environment while the other compartment remained empty. In this context, we identified a behavioral control event called Gap Surrogate. The initiation of this event was defined when the animal entered the region close to the gap with its nose, and its conclusion was marked in the third frame when the animal exited the gap. To form Surrogate Dyads, habituation recordings of individual dyad members, within which the Gap Surrogate events were identified, were artificially merged. This merging simulates social interaction events. By generating Surrogate Dyads, we established a control condition closely resembling the experimental context of the interacting dyads but without the presence of a conspecific on the other side of the barrier. When studying interactions between two-time series, distinguishing true coupling or correlation from random or spurious connections is crucial. Surrogate data help establish a null hypothesis by creating artificial datasets with statistical properties similar to the original but, in our context, without the presumed interaction. This is particularly important when dealing with oscillations at nearby frequencies produced by different mechanisms. Using surrogates that retain the original power spectrum helps prevent false positives in detecting coupling. Furthermore, these dyads facilitate the correction of artifact phenomena that could be generated by external sources or naturally present (e.g., simultaneous switching on of the LED of the MiCe-μScope or physiological phenomena related to cardiovascular or respiratory activity). These events are present in all recordings, regardless of the behavioral repertoire exhibited by the animals^75^.

After reviewing all the behavior videos, we used the BORIS software to download a “.csv” file. This file contained a data frame with the segmentation data, including the start and end times of the three behavioral events, categorized by animal and session. The generated data frame was subsequently used for the calcium imaging analysis to correlate the behavior and cortical activity.

### Calcium imaging analysis

With the MiCe-μScope, we captured cortical activity distributed across the entire cortex. Utilizing the Thy1-GCaMP6f transgenic line, we recorded neuronal responses expressed as fluorescence fluctuation, serving as an indirect indicator of calcium activity. Images, with gray levels representing the fluorescent light captured by the MiCe-μScope, are acquired as image stacks and saved in “*.tif*” format. Before proceeding with the chosen method for data analysis, a series of preprocessing steps are typically undertaken. Primarily, we align the image stacks across sessions and animals using anatomical landmarks marked during surgery. Subsequently, we normalize the fluorescence signal as part of the preprocessing procedure.

#### Image alignment

Image alignment is crucial for comparing neural activity in specific cortical regions across recordings from different sessions, days, and animals of the dyads. This step is also essential for parceling the cortex into regions according to a reference atlas^76^. To perform image registration, we utilize a custom-made Python tool based on the PyQtGraph library and the Allen atlas Python API (Allen SDK). This tool employs an affine transformation, estimated by mapping the coordinates of bregma and lambda to those of the Allen Brain Atlas, to register images. During the alignment process, bregma and lambda, which generally have an anatomical distance of 4.2 mm, serve as reference points for applying the parcellation mask to the cortex’s image plane. Once the images are registered, the parcellation mask can be applied to segment the entire cortex into different regions.

#### ΔF/F_0_ calculation

Normalization of the raw fluorescence signal is essential to compensate for intrinsic variability in the measurement system. This variability can arise from fluctuations in LED illumination intensity, spatial non-uniformity in illumination, or variations in the expression of the fluorescent indicator in the cortical layer of the animal. To account for these factors and ensure comparable values among animals on different sessions and days, the fluorescence signal is expressed as *ΔF/F*. This value is calculated using the following formula: 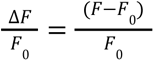,where F is the raw value of the fluorescence signal for the given pixel, and F_0_ represents the reference fluorescence, which is the mean fluorescence per pixel over time. Generally, this ratio is reported as a percentage.

#### Wavelet transform coherence analysis

To assess the coherence between two signals, the Wavelet Transform Coherence (WTC) is employed. This technique facilitates time coupling through variable-length windows, accommodating short windows for high frequencies and longer windows for low frequencies. This enables the measurement of cross-correlation between signals in the time-frequency space, revealing coherent phase relationships that may imply causality or a potential link between the two-time series. WTC excels in detecting locally locked-in phase events. The output of this analysis is a correlation coefficient in both time and the space of frequencies.

The calcium signal extracted was utilized to gauge inter-brain coupling in various behavioral conditions (Gap Interacting, Gap Non-Interacting, and Gap Surrogate). For studying and quantifying the coupling between these signals, WTC analysis was applied. This technique offers the advantage of measuring synchrony levels between two-time series concerning time and frequency. WCT, with its ability to unveil locally locked-in phase events not easily discernible with Fourier Transform, coupled with its adaptability to non-stationary biomedical signals by avoiding the need for a fixed window, makes it a robust analytical tool.

In this study, Morlet’s wavelet was selected as the mother function for its balanced characteristics in temporal localization and frequency. Morlet’s wavelet is multiplied by the signal (calcium profile), and the integration is performed at each instant. As the mother wavelet slides over the time series, higher product values indicate more similar profiles, while dissimilar points result in a product of zero. WCT reflects the relationship between the mother wavelet, at varying levels of compression and dilation (scaling factor), and the time series. The resulting spectrum illustrates time on the x-axis, representing the entire 310-second recording of neuronal activity, while the y-axis displays the frequencies of the calcium profile related to cortical activity. Considering the kinetics of the GCaMP6f indicator, a sampling frequency of 25 Hz was chosen to effectively capture calcium transients. With this sampling frequency, the maximum analyzable frequency resolution by WCT is half of it, equivalent to 12.5 Hz. The degree of synchrony between signals is portrayed on a colorimetric scale by coherence, ranging from 0 to 1. A coherence of 1 signifies perfect synchrony, while 0 indicates the absence of coherence. This analysis was conducted for all acquisitions, both during Solitary and Interaction sessions. Utilizing the behavior segmentation facilitated by BORIS and the obtained Data Frame, time windows corresponding to Gap Interacting, Gap Non-Interacting, and Gap Solitary events were extracted from the complete spectrum.

### Statistical analysis

All statistical analyses for this study were performed with custom Python scripts. All data are reported as mean ± SD unless otherwise indicated. In the cat plot, the error bars represent the 95% confidence interval. In the line plot of % Δ*F*/*F*_0_ the coherence spectrum of the entire cortex, the highlighted line represents the mean and the shadow represents the 95% confidence interval (Figures 1D, 1F, 3A, and 3B). Mixed-effect ANOVA was used for behavioral data analysis to evaluate the time spent in the ROI (center and outer) (Figure 1I) and the distance traveled during Interacting and Solitary sessions (Figure 2E). Welch’s t-test was used to examine the total distance traveled, still time, and speed in the two experimental groups (MiCe-μScope and Control mice) (Figures 1J, 1K, 1L). Repeated-measures ANOVA was used to examine effects on duration and number of nose poke events (Figures 2H and 2I). Repeated-measures ANOVA was also used to analyze the effect of maximum coherence peaks, divided by frequency bands, modulated by behavioral conditions. In this case, the Holm-Bonferroni correction was applied for the post-hoc t-test (Figures 3C, 3D, 3E, 3F). Paired t-test was used to compare the difference in duration of the first interaction compared to the subsequent ones (Figure 2J). A p-value of 0.05 was used as the criterion for a significant statistical difference. No statistical methods were used to predetermine sample sizes. All statistical analyses and related figures were created using Python and were assembled in Inkscape.

## ACKNOWLEDGMENTS

The author(s) declare financial support was received for the research, authorship, and/or publication of this article. This research was supported by the Italian National Recovery and Resilience Plan (NRRP), M4C2, funded by the European Union – NextGenerationEU Project Project IR0000023, CUP B53C22001810006, SEE-LIFE and Project COoperation and BRAin-Synchrony: a multiscale and translable approach CUP B53D23026560001.

---

**Figure S1.**
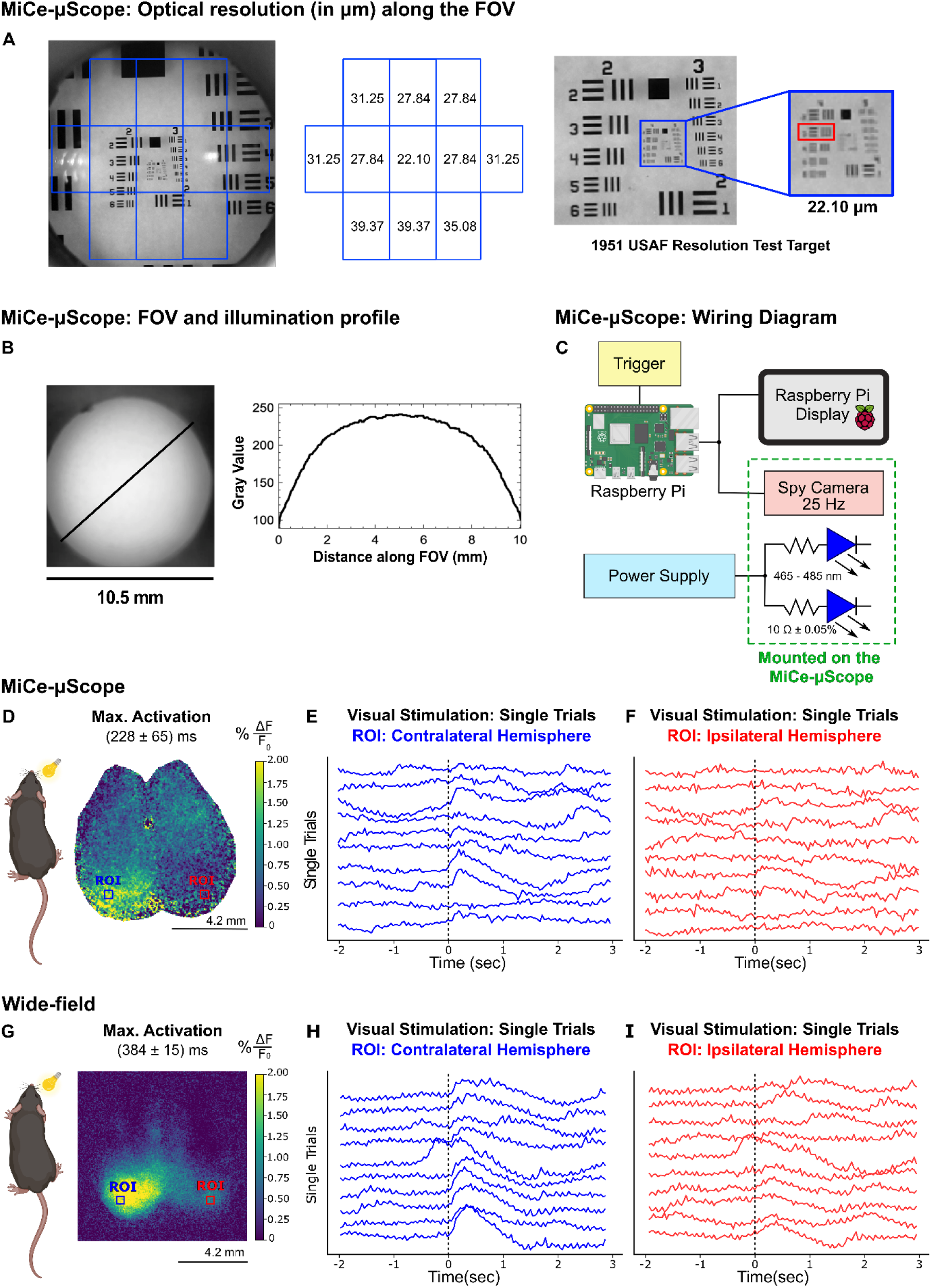
Optical characterization of the mice-μscope and benchmarking the mice-μscope with the conventional wide-field microscope at the single trial level. (A) Optical resolution in micrometers (μm) across the Field of View (FOV) was calculated using the 1951 USAF Resolution test target. The resolution was determined by locating the USAF in 11 different positions. An average resolution of (31.00 ± 5.01) μm was obtained across the entire FOV, with values expressed as the mean value ± standard deviation. (B) Image of the FOV and plot of the illumination profile across the FOV. (C) Wiring diagram illustrating the connection between the Raspberry Pi, Raspberry Pi display for viewing both hemispheres, and the Spy Camera for Raspberry Pi 5MP used for calcium imaging mounted on the top of the MiCe-μScop. The two blue LEDs (465 - 485 nm) are controlled by an external power supply. The components mounted on the MiCe-μScop are enclosed within the green dotted rectangle. (D and G) Maximal cortical activation frames: frames display maximal cortical activation following visual stimulation in the right eye. Blue and red ROIs indicate the contralateral and ipsilateral regions of the visual cortex from which the fluorescence signals were extracted with the MiCe-μScope (D) and wide-field microscope (G). (E-F and H-I) Individual calcium profiles: plot with individual calcium profiles depict the sensory responses evoked by individual visual stimulations in the visual cortex of the contralateral (blue) and ipsilateral (red) hemisphere, acquired with the MiCe-μScope (E-F) and wide-field microscope (H-I).

**Figure S2.**
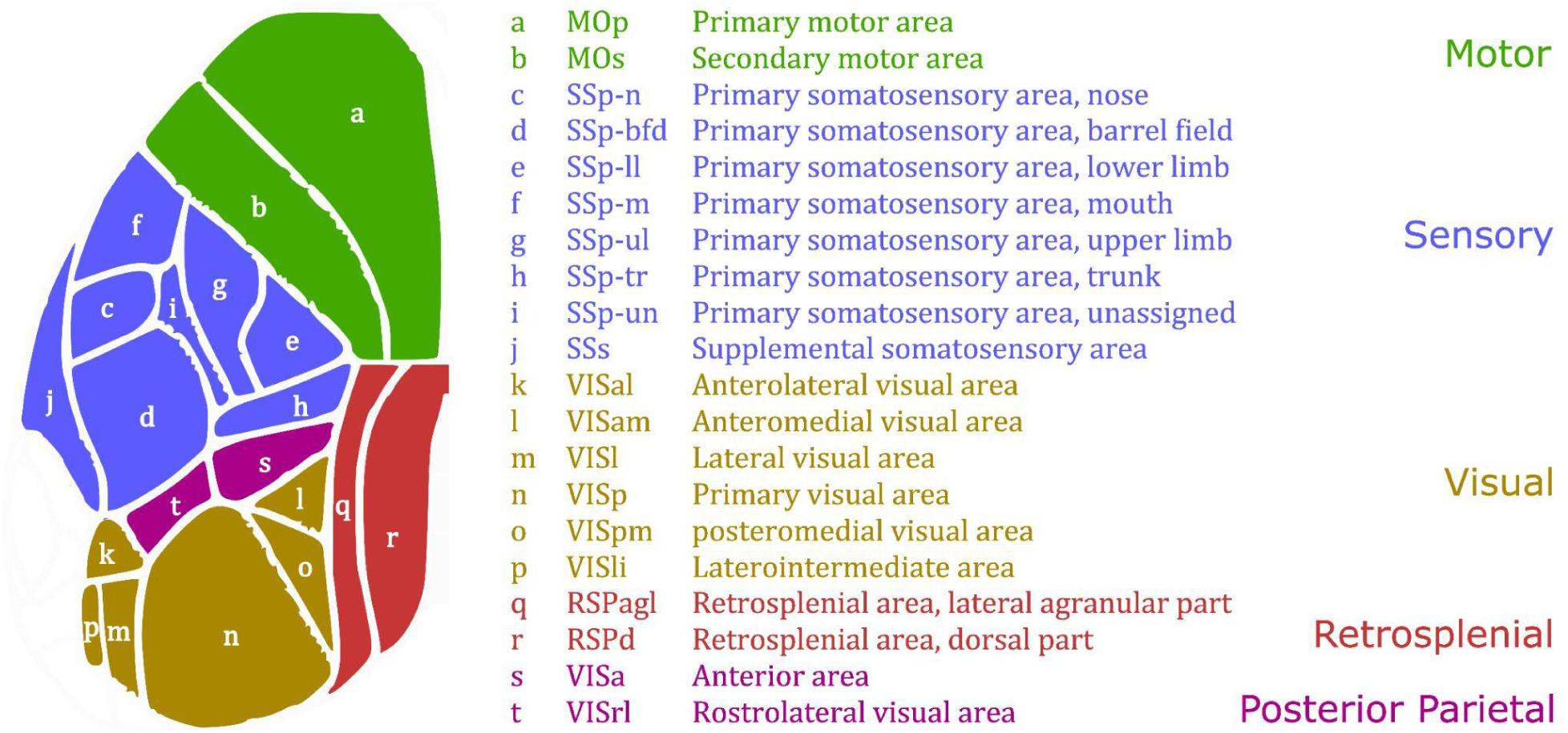
Mouse brain hemisphere with Allen map. Representation of a cerebral hemisphere with Allen’s Mouse Brain Atlas parcellation map aligned. Each letter corresponds to the acronym of the corresponding cortical area. The motor areas are grouped in green, sensory areas in blue, visual areas in brown, retrosplenial areas in red, and posterior parietal areas in purple.

